# Mining the human tonsillar microbiota as autoimmune modulator

**DOI:** 10.1101/719807

**Authors:** Jing Li, Shenghui Li, Jiayang Jin, Ruochun Guo, Xiaolin Sun, Jianping Guo, Fanlei Hu, Yanying Liu, Yuebo Jin, Yunshan Zhou, Wenjing Xiao, Yan Zhong, Fei Huang, Hudan Pan, Rentao Yang, Yuanjie Zhou, Kaifeng Deng, Lijun Wu, Liang Liu, Junjie Qin, Jun Wang, Jing He, Zhanguo Li

**Author notes:** Corresponding authors Prof. Zhanguo Li, Department of Rheumatology and Immunology, Peking University People’s Hospital, Beijing, 100044, China;. Prof. Jing He, Department of Rheumatology and Immunology, Peking University People’s Hospital, Beijing, 100044, China;. Prof. Jun Wang, CAS Key Laboratory for Pathogenic Microbiology and Immunology, Institute of Microbiology, Chinese Academy of Sciences, Beijing, 100101; Dr. Junjie Qin, Promegene Translational Research Institute, Shenzhen, 518110, China;. These authors contributed equally to this work (Jing Li, Shenghui Li, Jiayang Jin).

## Abstract

Palatine tonsils are important lymphoid organs featuring constant cross-talks between the commensal microorganisms and immune system, and have been implicated as critical autoimmunity origins for immune-related diseases, including rheumatoid arthritis (RA), a common autoimmune disorder. However, there was no evidence to show link between tonsillar microbiota and RA. Here, we identified a significant dysbiosis of RA tonsillar microbiota, with loss of *Streptococcus salivarius* and its functional molecules salivaricins (a type of antibacterial peptides). Consistent with the niche-preference, *S. salivarius* and salivaricins administrated intranasally or intraorally conferred prophylactic and therapeutic efficacies against experimental arthritis. Moreover, we demonstrated, for the first time, that *S. salivarius* and salivaricins exerted immunosuppressive capacities via inhibiting CD4^+^effector T cell subsets and autoantibody production in mice and human. These results uncover a communication between tonsillar microbiota and host autoimmunity, and identify the active components from tonsillar microbes in modulating immune homeostasis.

**One sentence summary:** Tonsillar microbiota regulate host autoimmunity via antibacterial peptides

---

Human palatine tonsils represent mucosa-associated lymphoid organs that serve as the central handing sites against microbial antigens. The tonsils undergo continuous immune responses that can exert impact on distant organs (*1, 2*). Previous studies have highlighted the important roles of tonsils in the pathogenesis of systemic diseases (*3*), including rheumatoid arthritis (RA) (*4*), a prototypical autoimmune disease with high morbidity and disability rate (*5*). The tonsils have been implicated as autoimmunity origins for RA, as evidenced by the findings that simultaneous and clonally identical T cell expansion was detected in the tonsils and synovium of the same RA patients (*4, 6*). This idea was further supported by the observations that tonsil-derived mononuclear cells could migrate to the synovial tissues in mice (*7*). However, the underlying mechanisms by which the tonsils contribute to RA development remain poorly defined.

Emerging evidences have emphasized the critical roles of microbiota and mucosal immune system in the etiopathogenesis of autoimmune disease (*8, 9*). Previous reports have demonstrated the indispensable participation of microorganisms in experimental arthritis (*10, 11*). Recent studies provided further evidence for the involvement of gut and lung microbiota in RA (*12, 13*). The tonsillar microbiota is in intimate contact with crucial antigen-handling lymphoid tissues, and alterations of human-microbiota mutualism in the tonsil can trigger local infections or systemic diseases (*14, 15*). However, no report has elucidated the direct link between tonsillar microbiota and autoimmune disorders. Here, we revealed a communication between tonsillar microbiota and host autoimmunity, and identified key tonsillar microbes and corresponding functional molecules in regulating immune homeostasis.

To profile the tonsillar microbiota, we began with 16S rRNA gene sequencing on tonsillar swap samples from 121 RA patients and 99 healthy controls (Table S1). The tonsil harbored a highly diversified microbial consortium with only a minimal proportion of microbial taxa shared with gut microbiota (Table S2 and fig. S1). Tonsillar microbiota of RA patients displayed similar within-sample (α) diversity, yet significant increase in between-sample (β) diversity compared to healthy controls (fig. S2). Principle coordinate analysis revealed a significant separation of RA samples from controls (Adonis test, *P*<0.001, Fig. 1A). Other confounding variables, including sex, age, body mass index, and dietary patterns, explained smaller proportions of variation of the tonsillar microbiota (Table S3). Additionally, the composition of RA tonsillar microbiota was correlated with disease activity and medication treatment (Table S3), especially that medication could reverse to a certain extent the microbial differences (fig. S3).

**Fig. 1.**
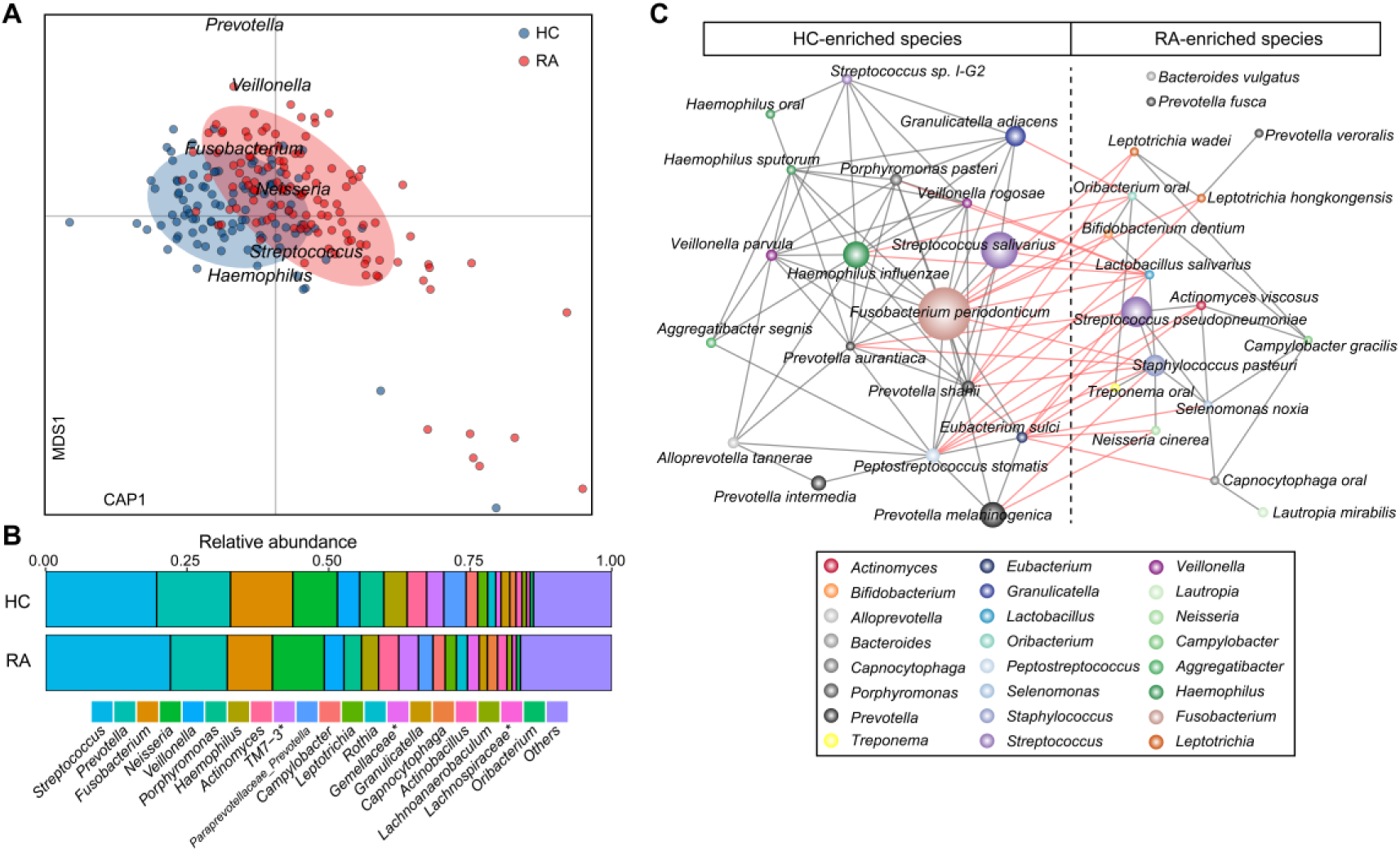
Difference of tonsillar microbiome between RA patients and healthy controls. **(A)** Distance-based redundancy analysis (dbRDA) based on Bray-Curtis dissimilarity between microbial genera, revealing a RA-associated microbial dysbiosis compared to controls. We showed the primary constrained axis (CAP1) and primary multidimensional scaling (MDS1), while top six genera contributors are plotted by their loadings in these two axes. RA, rheumatoid arthritis, n=121; HC, healthy controls, n=99. Adonis test, *P* < 0.001. **(B)** Composition of the tonsillar microbiota at genus level. Unclassified genera under a higher rank were marked by asterisks. **(C)** Correlation network between 35 RA-associated species that were significantly different between healthy controls and RA patients. Red and grey edges respectively denoted negative and positive correlation (Spearman’s correlation coefficient |ρ| > 0.35, *q*<0.05). Species of same genera were noted by identical colors.

With respect to specific taxonomical groups, *Prevotella, Fusobacterium* and *Haemophilus* were significantly depleted in RA patients, whereas *Streptococcus* was enriched in patients but not significant (Fig. 1B and fig. S4). We further assigned the operational taxonomical units (OTUs) into closest species, and identified 35 species differentially abundant in RA patients and healthy controls (Fig. 1C and Table S4). Notably, patterns of these RA-related signatures could be partially reversed by RA medication (fig. S5), implicating the tonsillar microbiota as potential derivers in RA. Of the two major members of *Streptococcus, S. salivarius* was significantly reduced in RA patients, especially in treatment-naïve patients, whereas *S. pseudopneumoniae* was enriched in RA microbiota (Fig. 1C and fig. S5). *Lactobacillus salivarius*, a recently-reported species that over-represented in multiple body sites of RA patients (*12*), showed an increase in RA tonsil as well. Interestingly, *S. salivarius* and *L. salivarius* exerted antagonistic relationship with each other (Fig. 1C). Finally, random forest analysis yielded a high performance model (area under the curve = 0.93) for discriminating RA patients from healthy controls (fig S6).

To determine the functional consequences of taxonomic differences, we next performed whole-metagenome sequencing on tonsillar microbiota from 32 RA patients and 30 healthy controls (Table S5). We greatly extended the data depth of tonsillar microbiota that was not well-focused on in Human Microbiome Project (*16*). Expectedly, the RA-associated taxonomic signatures in 16S sequencing analyses were validated in the metagenomic data with concordant tendency (Table S6). Also, the functional profiles of RA and control microbiomes were remarkably differed (Adonis *P* < 0.001; fig. S7), and 78 of the 504 microbiome functional modules were significantly associated to RA status (false discovery rate < 0.1; Table S7). Of which, the 72 RA-reduced modules were commonly involved to metabolism of various organics, transport systems of some small molecules or ions, and two-component regulatory system, whereas the 6 RA-enriched enzymes encoded the major modules of methanogenesis. Reducing of compounds metabolism and transport systems agree well with the observations in the gut and oral microbiomes of RA patients (*12*) (Table S7). Moreover, species depleted in RA were frequently correlated with these functional modules lower in RA (Fig. 2A and fig. S8), and especially, positively correlated to richness indexes (Fig. 2A), suggesting that the lack of some key species and corresponding functions might be an important factor in RA pathogenicity.

**Fig. 2.**
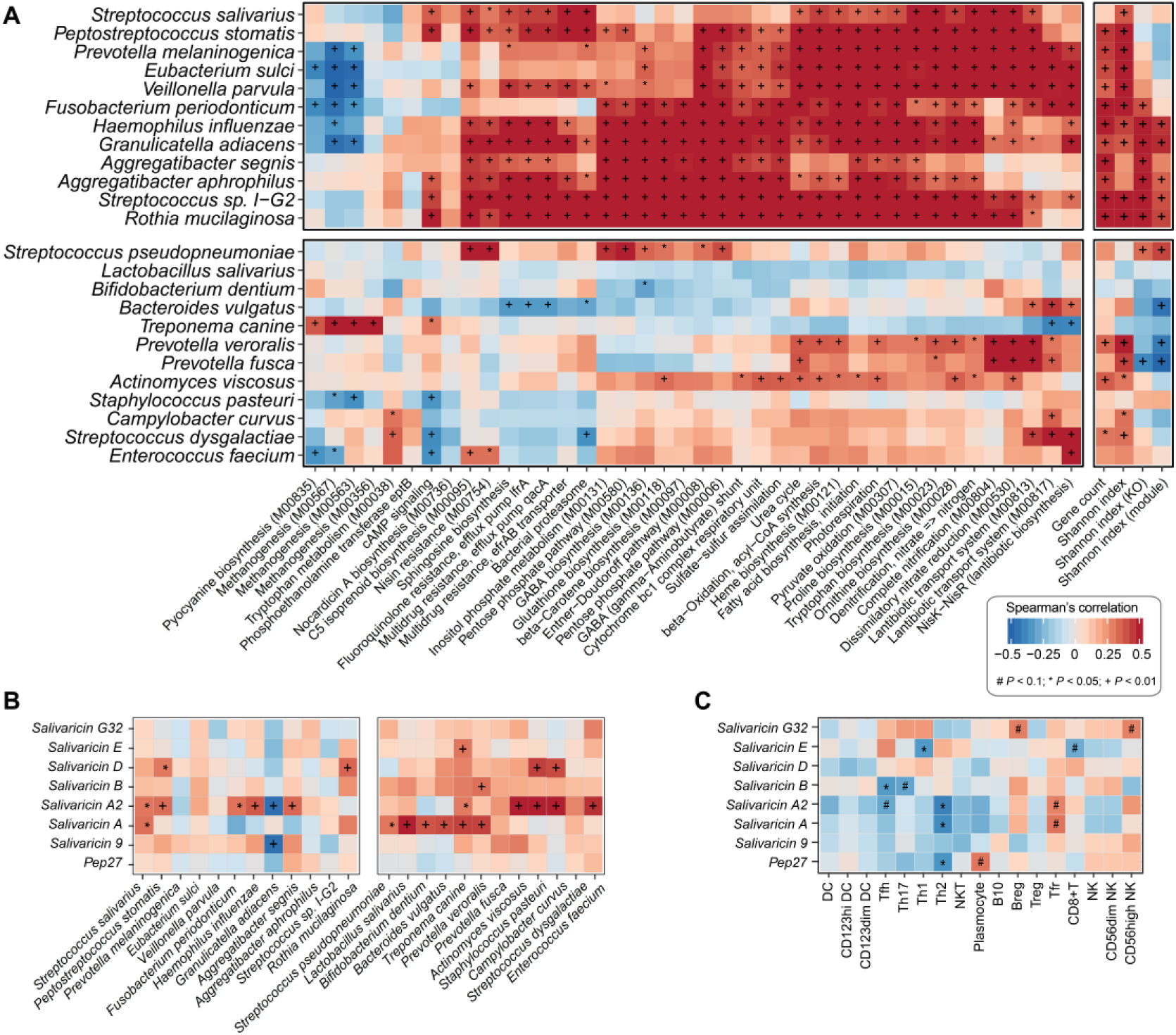
Lantibiotics as key differentiated microbial functional molecules in metagenomic analysis. **(A)** Heatmap showing the Spearman correlations of RA-associated species with functional modules (left panel) and microbial community diversity (right panel) in tonsillar microbiota. (**B and C)** Heatmap displaying associations of lantibiotics genes from tonsillar microbiota with RA-associated species (B) and circulating immune cells (C). DC, dendritic cells; Th, T helper cells; Tfh, follicular helper T cells; Tfr, follicular regulatory T cells; B10, interleukin-10 producing B cells; Breg, regulatory B cells; Treg, regulatory T cells; NK, natural killer; NKT, natural killer T cells. Significance determined using Spearman correlation analysis, n=62, ^#^*P* < 0.1, \**P* < 0.05, +*P* < 0.01.

We paid specific attention to the biosynthesis and transport modules of lantibiotics that deficient in RA tonsil. Produced by various bacteria and exhibit antimicrobial properties, lantibiotics play crucial roles in maintaining microecological homeostasis (*17*). It has been suggested that Streptococci especially *S. salivarius*, commonly produce lantibiotics as putative anti-competitor or signaling molecules (*18*). We identified 245 genes encoding lantibiotics in our metagenomic data and they were mainly involved in the biosynthesis of salivaricins produced by *S. salivarius* and Pep27 synthesized by *S. pneumoniae* (Table S8). Notably, four salivaricins, A2, B, A, and G32, were more abundant in the control cohort (Table S8), and positively correlated with *S. salivarius* and other RA-reduced species (Fig. 2B). Of those, salivaricin A2 and B demonstrated highest genetic diversity and abundances, particularly in healthy controls (Table S8 and fig. S9).

We further investigated the correlations of microorganisms with systemic immune responses. A number of RA-related tonsillar microbes showed associations with host circulating immune cells and autoantibody (Fig. 2C, fig. S10, and Table S9). To our most interest, the loss of *S. salivarius* and genes encoding salivaricins correlated with elevated titers of anti-cyclic citrullinated peptide (CCP) antibody and increased proportions of precursor follicular helper T (Tfh) and T helper 17 (Th17) cells, two key pathogenic immune mediators in RA (*19, 20*). Together, these data implied that tonsillar microbiota might take active part in the autoimmune responses of RA.

To directly test the immunomodulatory effects of *S. salivarius*, we performed *in vitro* experiments using peripheral blood mononuclear cells (PBMCs) isolated from healthy individuals. As expected, *S. salivarius* strain K12, a safe and well tolerated oral probiotic (*18*), could negatively regulate cytokine responses of PBMCs co-incubated with T cell-specific stimulus anti-CD3 plus CD28 antibody (fig. S11). To further apply the ability of *S. salivarius* in suppressing immune-mediated pathological processes *in vivo*, we introduced this species to collagen-induced arthritis (CIA) model, a well-characterized murine model of arthritis resembling human RA (*21*). In the preventative studies, *S. salivarius* K12 inoculated intra-nasally, intra-orally or combined administration all significantly decreased incidence and severity of arthritis without notable effects in body weight (Fig. 3, A and B; fig. S12, A and B). In contrast, *L. salivarius* administrated intra-nasally and -orally exhibited no apparent protection from CIA (fig. S12, C and D). Additionally, *S. salivarius* K12 supplemented by intragastric gavage showed no protective efficacy against CIA either (fig. S12, E and F), suggesting niche-preferred properties of RA-related tonsillar microbe in modulating arthritis. For CIA model with already-developed arthritis, *S. salivarius* K12-treated mice also showed significantly relieved disease severity and reduced inflammatory cell infiltration in the joints (fig. S12, G and H), supporting a clinical applicability of *S. salivarius* K12 in treating RA.

**Fig. 3.**
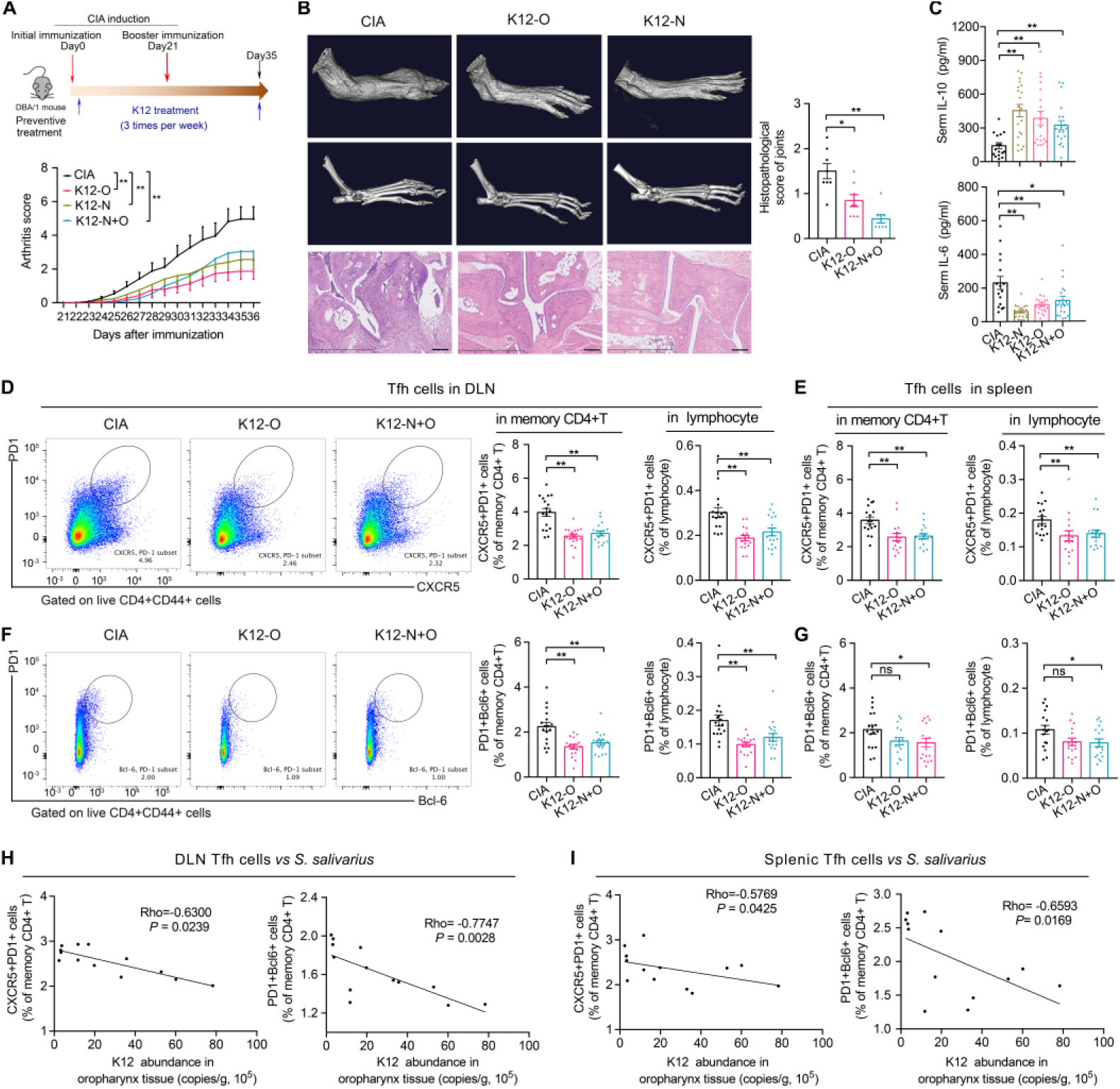
*S. salivarius* suppresses immune responses and protects against experimental arthritis. **(A)** The study protocol of a preventive regimen with a CIA mouse model (upper panel) and clinical arthritis scores in CIA mice with or without *S. salivarius* K12 (1*10^8^cfu/mice) inoculation. CIA, collagen-induced arthritis, n=26; K12-O, K12-inoculated intra-orally, n=25; K12-N, K12-inoculated intra-nasally, n=30; K12-N+O, K12-inoculated intra-orally and intra-nasally, n=30. cfu: colony forming unit. **(B)** Representative images of Mirco-CT and H&E-stained sections as well as histopathological scoring of the paws. Scale bars, 200 μm. n=8 per group. **(C)** Concentrations of serum IL-10 and IL-6. CIA, n=16; K12-O, n=20; K12-N, n=20; K12-N+O, n=20. (**D-G)** Representative flow cytometry plots with graph of frequency of CD4^+^CD44^+^CXCR5^+^PD1^+^Bcl6^+^ Tfh cells in draining lymph modes (DLN) and spleen of indicated groups. n=17 per group. **(H and I)** The scatter plots depict associations of frequencies of Tfh cells with *S. salivarius* K12 abundance in the oropharyngeal mucosal of mice, n=13. Data were pooled from two independent experiments and expressed as mean ± SEM. Significance determined using two-way ANOVA followed by Tukey’s multiple comparisons test (A), Mann-Whitney U test (B-G) or Spearman correlation analysis (H and I). \**P* < 0.05, \*\**P* < 0.01, ns: non-significant.

We then determined the immune effector molecules responsible for the anti-arthritis effectiveness of *S. salivarius* K12 by examining cytokines and lymphocyte subsets that mechanistically linked to the severity of CIA. Increased serum level of anti-inflammatory mediator IL-10 and decreased concentration of pro-inflammatory cytokine IL-6 (Fig. 3C) were detected in *S. salivarius* K12-treated mice. Furthermore, *S. salivarius* K12 markedly decreased the frequencies of Tfh cells in both draining lymph nodes (DLN) and the spleen (Fig. 3, D and E). Accordingly, the expression of Bcl-6, a transcription factor essential for the differentiation and function of Tfh cells (*22*), was also significantly reduced (Fig. 3, F and G). In addition, the expression and differentiation of Th17 cells were inhibited, while the proportions of Th1 and Treg cells were unchanged (fig. S13), suggesting selective modulations of *S. salivarius* K12 in CD4^+^ T cell subsets. More importantly, these reduced systemic immune responses could be linked to *S. salivarius* K12 colonization in the oropharyngeal mucosal of CIA mice, as revealed by the negative associations of Tfh cell frequencies and the positive correlation of serum IL-10 level with *S. salivarius* K12 abundance (Fig. 3, H and I; figs. S14 and S15), confirming the active immunosuppressive effect of *S. salivarius* K12 in protecting against arthritis progression.

Commensal microbes calibrate immune responses in large part by producing small molecules that mediate host-microbial interactions (*23, 24*). Thus, we explored the immunomodulatory properties of salivaricins, the key anti-competitor or signaling molecules mediating the benefits of *S. salivarius (18)*. Similar with *S. salivarius* K12, salivaricin A2 and B (purified from *S. salivarius* K12 culture medium, fig. S16) exerted immunosuppressive effects in cultured human PBMCs under T cell-specific stimulus, as shown by the reduced levels of IL-21, IL-17A, and anti-immunoglobulin (Ig) G antibody (fig. S17, A-G). Moreover, salivaricin A2 and B could significantly down-regulated the relative percentages of precursor Tfh cells, a primary producer of IL-21 and a key promotor to antibody production (*22*), but had no detectable effect on Treg cells (fig. S17, H and I).

To further test the anti-arthritis efficacy of salivaricins, we directly administrated chemosynthetic salivaricin A2 or B intra-orally to CIA mice. The chemical synthesized salivaricin A2 and B (fig. S18) possessed similar antibacterial (Table S10) (*25, 26*) and immunoregulatory activities (fig. S19) as to the bacteria-produced salivaricins. In the preventative regimen, salivaricin A2 or B supplemented mice exhibited evident lower CIA incidence and disease scores with no body weight changes compared to the vehicle-treated mice, both clinically and histologically (Fig. 4, A-C and fig. S20, A-D). Furthermore, salivaricin A2 or B treated mice showed reduced titer of serum autoantibody anti-collagen type II (CII) and increased concentration of serum IL-10 with no notable changes for IL-6 or IFN-γ (Fig. 4, D and E; fig. S20, E and F). In addition, salivaricin A2 and B exerted similar modulations in lymphocyte subsets with that of *S. salivarius* K12, as evidenced by remarkably reduced frequencies and differentiation of Tfh and Th17 cells as well as decreased percentages of plasmablast in both DLN and the spleen (Fig. 4, F-K and fig. S21). For the established CIA model, salivaricin A2 and B also showed therapeutic efficacies in ameliorating disease severity and reducing autoimmune responses (fig. S22). Together, these results demonstrated that the protective mechanisms of *S. salivarius* K12 in experimental arthritis might be majorly dependent on the role of salivaricins.

**Fig. 4.**
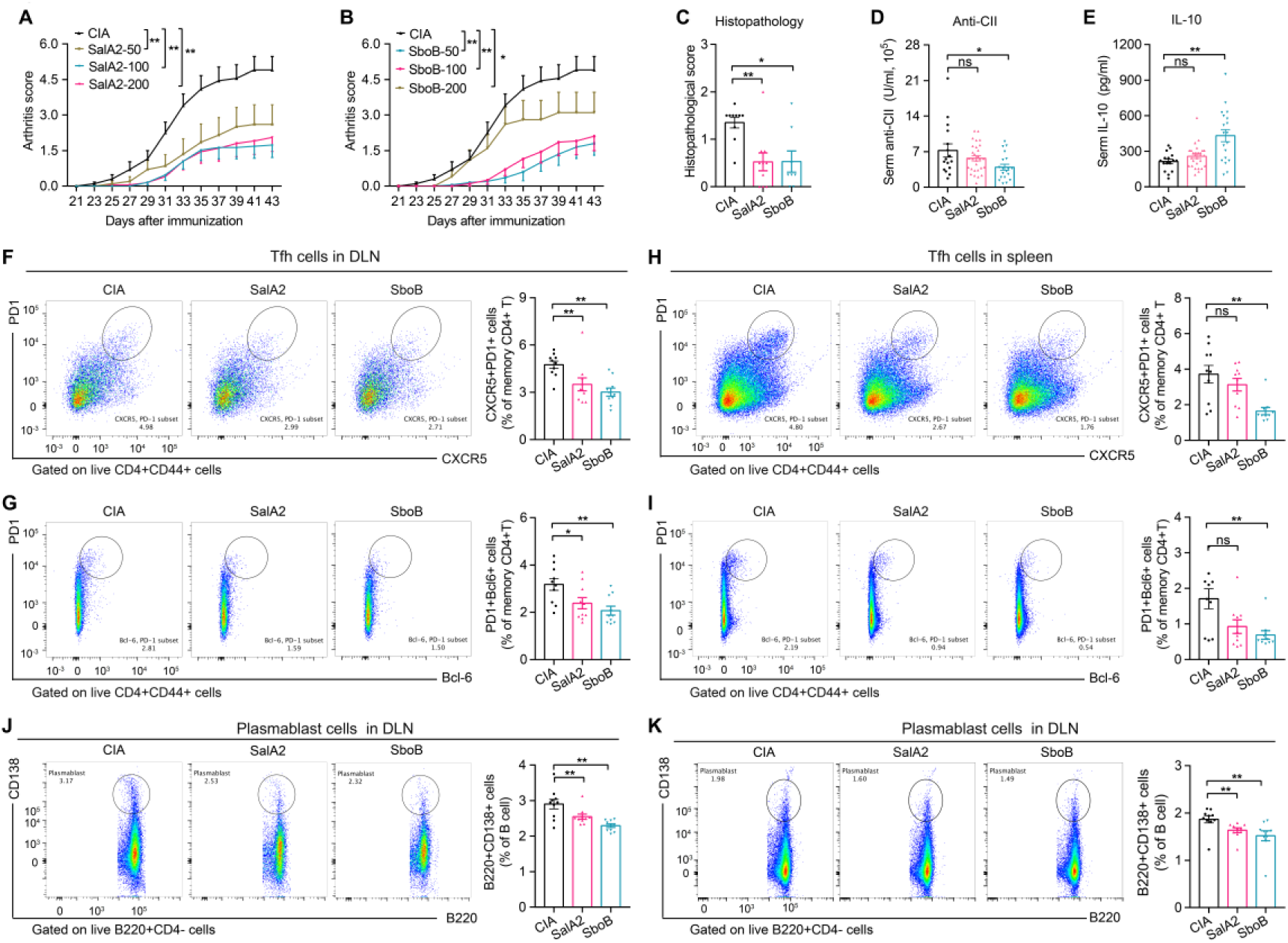
Salivaricins recapture the effect of *S. salivarius* in immune regulation and arthritis prevention. **(A and B)** Clinical arthritis scores in CIA mice with or without salivaricin A2 (A) or salivaricin B (B) intro-orally administration. SalA2: salivaricin A2, SobB: salivaricin B, n=20 for each group except for SobB-200 (n=10). Unit: μg/mice. **(C)** Histopathological scoring of the paws. CIA, n=10; SalA2, n=10; SboB, n=8. **(D and E)** Serum concentrations of anti-collagen type II (CII) antibody and IL-10. CIA, n=16; SalA2, n=29; SboB, n=19. **(F-K)** Representative flow cytometry plots with graph of frequencies of CD4^+^CD44^+^CXCR5^+^PD1^+^Bcl6^+^Tfh cells and B200^+^CD138^+^plasmablast in DLN and the spleen of indicated groups. n=10/group. Data were pooled from two independent experiments and expressed as mean ± SEM. Significance determined using two-way ANOVA followed by Tukey’s multiple comparisons test (A and B) or Mann-Whitney U test (C-K), \**P* < 0.05, \*\**P* < 0.01, ns: non-significant.

In summary, the key finding of the present study is that the dysbiotic tonsillar microbiota contributes to autoimmune disorders via the missing production of important bacterial peptides including salivaricins produced by *S. salivarius*, thus losing the ability to properly modulate CD4^+^ effector T cell subsets and autoantibody production, which could act as signals sent out by the tonsils to regulate autoimmune response at distal sites. Our work substantially identifies an interesting communication between tonsillar microbiota and autoimmune diseases, and provides strategies in which specific tonsillar bacterium maintain immune homeostasis.

